# A CLEAR pipeline for direct comparison of circular and linear RNA expression

**DOI:** 10.1101/668657

**Authors:** Xu-Kai Ma, Meng-Ran Wang, Chu-Xiao Liu, Rui Dong, Gordon G. Carmichael, Ling-Ling Chen, Li Yang

## Abstract

Sequences of circular RNAs (circRNAs) produced from back-splicing of exon(s) completely overlap with sequences from cognate linear RNAs transcribed from the same gene loci with the exception of their back-splicing junction (BSJ) sites. Examination of global circRNA expression from RNA-seq datasets generally relies on the detection of RNA-seq fragments spanning BSJ sites, but a direct comparison of circular and linear RNA expression from the same gene loci in a genome-wide manner has remained challenging. This is because quantification of BSJ fragments differs from that of linear RNA expression that uses normalized RNA-seq fragments mapped to the whole gene bodies. Here, we have developed a computational pipeline for <u>c</u>ircular and linear RNA <u>e</u>xpression <u>a</u>nalysis from ribosomal-RNA depleted RNA-seq (CLEAR, https://github.com/YangLab/CLEAR). A new quantitation parameter, FPB (fragments <u>p</u>er <u>b</u>illion mapped bases), is applied to evaluate circular and linear RNA expression individually by fragments mapped to circRNA-specific BSJ sites or to linear RNA-specific splicing junction (SJ) sites. Then, circular and linear RNA expression are directly compared by dividing FPB_circ_ by FPB_linear_ to generate a CIRCscore, which indicates the relative circRNA expression using linear RNA expression as the background. Highly-expressed circRNAs with low cognate linear RNA expression background can be identified for further investigation.

## INTRODUCTION

Eukaryotic pre-mRNA splicing is catalyzed by spliceosomes to join upstream 5’ splice donor sites with downstream 3’ splice acceptor sites to produce linear (m)RNAs. Downstream 5’ splice donor sites can also be linked to upstream 3’ splice acceptor sites, referred to as back-splicing, leading to the production of circular RNAs (circRNAs) from thousands of gene loci (1-3). Unlike most mature linear RNAs (including both coding and long noncoding RNAs), circRNAs lack 3’-end poly(A) tails, resulting in their depletion in poly(A)+ RNA-seq datasets. By taking advantage of analyzing RNA-seq datasets that profile non-polyadenylated transcripts, including ribosomal RNA depletion (ribo–) RNA-seq, and computational approaches that aim to identify fragments mapped to BSJ sites (4,5), a large number of circRNAs have been successfully profiled as being co-expressed with their cognate linear RNAs from the same gene loci (2,3,6-8). Recent research has shown that the biogenesis of circRNAs is catalyzed by canonical spliceosomal machinery and modulated by both *cis*-elements and *trans*-factors (1-3,9, 10). Importantly, increasing lines of evidence have revealed that circRNAs are involved in physiological and pathological conditions with different modes of actions (6,11-14).

Despite these findings, comprehensive characterization of circRNA biogenesis and function has been impeded by the facts that the majority of circRNAs are processed from middle exons of genes and that their sequences almost completely overlap with those of their cognate linear RNAs except for the BSJ sites (2). No genome-wide method has been available to compare the expression of circRNAs with their cognate linear RNAs directly from RNA-seq datasets. The primary obstacle for direct expression comparison is owing to distinct strategies for circular and linear RNA quantification from mapped RNA-seq fragments. In general, RNA-seq fragments that are solely mapped to BSJ sites are used to represent circRNA expression, such as by raw or normalized (as known as FPM, fragments <u>p</u>er <u>m</u>illion mapped fragments, Figure 1A, left) fragment counts. On the other hand, RNA-seq fragments mapped to both exon bodies and exon-exon splicing junction (SJ) sites are summed up and normalized for linear RNA quantification, such as by FPKM (15) (fragments <u>p</u>er <u>k</u>ilobase of transcript per <u>m</u>illion mapped fragments, Figure 1A, right). Since FPM is unscaled to FPKM, the relative expression of most circRNAs is not comparable to their cognate linear RNAs when analyzing RNA-seq datasets.

**Figure 1.**
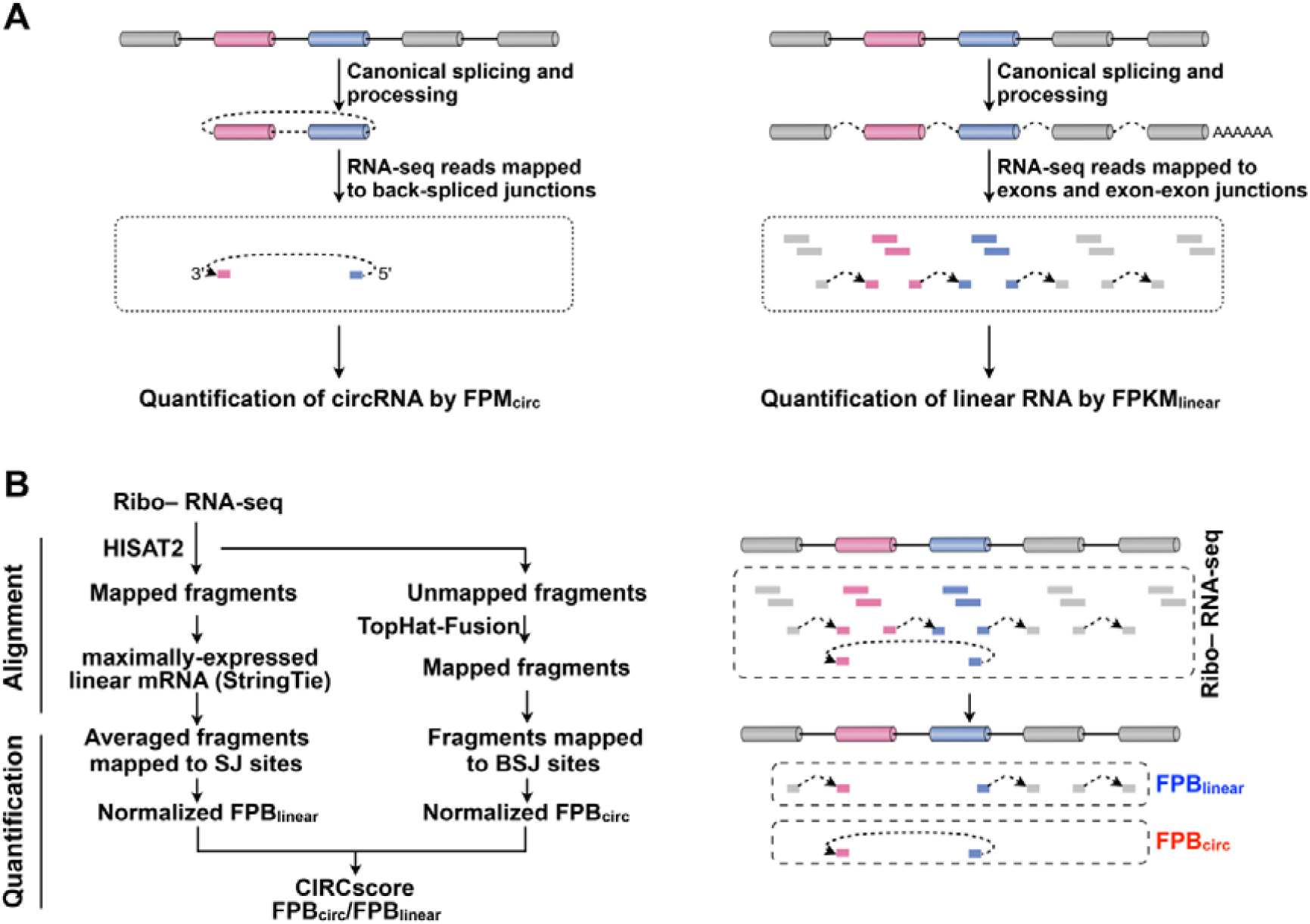
A computational pipeline for direct circular and linear RNA expression comparison. (**A**) Schematic drawing to show different quantification strategies for circRNAs (left) and linear RNAs (right). Thus, FPM (fragments <u>p</u>er <u>m</u>illion mapped fragments) for circRNA quantification and FPKM (fragments <u>p</u>er <u>k</u>ilobase of transcript per <u>m</u>illion mapped fragments) for linear RNA quantification are unscaled and incomparable. (**B**) Development of the computational pipeline for <u>c</u>ircular and linear RNA <u>e</u>xpression <u>a</u>nalysis from ribosomal-RNA depleted (ribo–) RNA-seq (CLEAR). Schematic diagram of the CLEAR pipeline (left) and the same strategy for circular or linear RNA quantification by splicing junction (SJ) or back-splicing junctions (BSJ) site mapped fragments <u>p</u>er <u>b</u>illion mapped bases (FPB). CIRCscore that divides FPB_circ_ by FPB_linear_ can be achieved for direct circular and linear RNA expression comparison.

To solve this problem, we have developed a computational pipeline for <u>c</u>ircular and linear RNA <u>e</u>xpression <u>a</u>nalysis from ribosomal-RNA depleted RNA-seq (CLEAR, Figure 1B). With the CLEAR pipeline, RNA-seq fragments mapped to circRNA-specific BSJ sites or linear RNA-specific SJ sites are individually normalized to evaluate circular or linear RNA expression, each in FPB (fragments <u>p</u>er <u>b</u>illion mapped bases). Unlike using the non-comparable FPM and FPKM values, circular and linear RNAs are both quantified by FPB values with the CLEAR pipeline, and thus can be directly compared by dividing FPB_circ_ by FPB_linear_ to generate a CIRCscore. In this scenario, relative circRNA expression can be evaluated by using linear RNA expression as an expression background, and highly-expressed circRNAs with low cognate linear RNA expression background can be identified for further functional studies.

## MATERIAL AND METHODS

### Direct circular and linear RNA expression comparison by the CLEAR pipeline

A computational pipeline for <u>c</u>ircular and linear RNA <u>e</u>xpression <u>a</u>nalysis from ribosomal-RNA depleted (ribo–) RNA-seq (CLEAR) was developed to achieve direct circular and linear RNA expression comparison. Ribo- RNA-seq datasets that profile both polyadenylated linear and non-polyadenylated circular RNAs in parallel are used for precise circular and linear RNA expression comparison.

The CLEAR pipeline includes two main steps: alignment and quantification (Figure 1B). For the alignment, ribo– RNA-seq fragments were first mapped by HISAT2 (version 2.0.5; parameters: hisat2 --no-softclip --score-min L,-16,0 --mp 7,7 --rfg 0,7 --rdg 0,7 --dta -k 1 --max-seeds 20) against GRCh38/hg38 human reference genome with known gene annotations (Supplementary Figure S1) for subsequent linear RNA quantification analysis. HISAT2-unmapped fragments were then mapped to the same GRCh38/hg38 reference genome using TopHat-Fusion (version 2.0.12; parameters: tophat2 --fusion-search --keep-fasta-order --bowtie1 --no-coverage-search) for subsequent circRNA quantification.

For the quantification, we developed a new FPB (fragments <u>p</u>er <u>b</u>illion mapped bases) value to quantitate linear RNA expression by HISAT2-mapped fragments to SJ sites of the maximally-expressed transcript annotation (Supplementary Figure S2). The maximally-expressed transcript of a given gene is selected with the highest FPKM value, which is calculated by StringTie (version 1.3.3; parameters: stringtie -e -G) from HISAT2 aligned BAM file (16). Fragments mapped to back-splicing junctions were retrieved from TopHat-Fusion as previously reported (version 2.3.6; parameters: CIRCexplorer2 parse -f -t TopHat-Fusion) (17) and normalized by totally-mapped bases to obtain FPB values for circRNA quantification.

Direct comparison of circular *vs* linear RNA expression can be achieved using the CIRCscore value that divides FBP_circ_ by FPB_linear_, which represents relative circRNA expression using linear RNA expression as the background.

### Availability and flexibility of the CLEAR pipeline

The CLEAR pipeline and its application can be downloaded from https://github.com/YangLab/CLEAR. Of note, other aligners, including TopHat2 (version 2.0.12; parameter: tophat2 -a 6 --microexon-search -m 2 -g 1) with known gene annotations (Supplementary Figure S3) or MapSplice (version 2.1.8 with default parameters) with gene annotations (ensGene_v89.txt updated at 2017/05/08) can also be used in the CLEAR pipeline with similar outputs.

In the CLEAR pipeline, comparable circular or linear RNA expression by FPBs and their direct comparison by the CIRCscore can be obtained directly from raw RNA-seq FASTQ files or processed RNA-seq results, such as CIRCexplorer2 output files (17). Please see https://github.com/YangLab/CLEAR for details.

### Cell culture

PA1 cells were purchased from the American Type Culture Collection (ATCC; http://www.atcc.org), and maintained in MEMα supplemented with 10% FBS, 1% Glutamine and 0.1% penicillin/streptomycin at 37°C in a 5% CO2 cell culture incubator. PA1 cells were routinely tested to exclude mycoplasma contamination.

### Comparison of FPB with qPCR quantification

Total RNAs from cultured PA1 cells were extracted with Trizol (Life technologies) according to the manufacturer’s protocol. Extracted RNAs were treated with DNase I (Ambion, DNA-freeTM kit), reversely transcribed with SuperScript III (Invitrogen) to produce cDNA and then applied for qPCR analysis. β-actin mRNA was examined as an internal control for normalization. Expression of examined linear and circular RNAs was determined from three independent experiments. The primers used are listed in Supplementary Table S1.

### RNA-seq datasets used in this study

Datasets used for this study include publicly available ribo–, poly(A)+, poly(A)–/ribo– and RNase R RNA-seq datasets from PA1 cell line (17,18), ribo– RNA-seq datasets of twelve tissues from ENCODE (19) (Supplementary Table S2), and ribo– RNA-seq datasets of 20 human hepatocellular carcinoma (HCC) samples and their paired control samples (GSE77509) (20).

## RESULTS

### Development of the CLEAR pipeline

The CLEAR pipeline was set up for direct circular and linear RNA expression comparison on a genome-wide scale (Figure 1B). Briefly, ribo– RNA-seq fragments are first mapped to the human reference genome/annotation (NCBI RefSeq genes, Supplementary Figure S1) by HISAT2 (21) to obtain fragments mapped to linear RNAs. Next, HISAT2-unmapped fragments are aligned by TopHat-Fusion (22) to retrieve fragments mapped to BSJ sites for circRNA quantification (Figure 1B, left, Alignment). Two characteristic features for circular and linear RNA quantification are applied in the CLEAR pipeline. Similar to circRNA quantification by RNA-seq fragments solely mapped to BSJ sites, fragments that only map to canonical splicing site (SJ) sites by HISAT2 are used for linear RNA quantification (Figure 1B, right). Different from commonly-used FPKM that counts fragments mapped to both exon bodies and SJ sites, linear RNA quantification by fragments only mapped to canonical SJ sites is comparable to circRNA quantification by those mapped to BSJ sites (Figure 1B). In addition, fragments mapped to SJ or BSJ sites are normalized by totally mapped bases, rather than by totally mapped fragments, to get FPB for linear or circular RNA quantification (Figure 1B, left). Direct circular and linear RNA expression comparison can be achieved with the CIRCscore that divides FPB_circ_ by FPB_linear_ (Figure 1B, left).

### Comparison of FPB with FPKM for linear RNA quantification

To evaluate the accuracy of FPB for RNA quantification, commonly-used FPKM values are obtained from the same HISAT2-mapped results. Basically, HISAT2-mapped results are first converted to BAM format by SAMtools (23). StringTie (16) is then used to calculate transcript expression by FPKM. Since multiple linear RNAs can be produced from a given gene locus, the average FPB value of fragments mapped to all SJ sites in the maximally-expressed linear transcript is used to represent the expression of this gene in the current study (Figure 1B and Supplementary Figure S2A, S2B).

With the requirement of FPB_linear_ > 0 and FPKM_linear_ > 0, linear RNA expression, when quantitated by FPB_linear_, is highly correlated with that by FPKM_linear_ in the PA1 cell line (18) (Figure 2A). Indeed, the value of FPB_linear_ is theoretically equivalent to that of FPKM_linear_ (Supplementary Figure S2C). Furthermore, FPB_linear_ is highly correlated with the expression level of thirteen linear RNAs measured by RT-qPCR in PA1 cells (Figure 2B, Supplementary Table S3). We observe a high correlation between FPB_linear_ and FPKM_linear_ when using different aligners, such as TopHat2 (24) and MapSplice (25), to analyze the ribo– RNA-seq dataset of PA1 (Supplementary Figure S3). Finally, FPB_linear_ is also highly correlated with FPKM_linear_ in ENCODE RNA-seq datasets from twelve examined human tissues (Supplementary Figure S4 and Supplementary Table S2). Collectively, these findings reveal that FPB_linear_ is applicable for linear RNA quantification.

**Figure 2.**
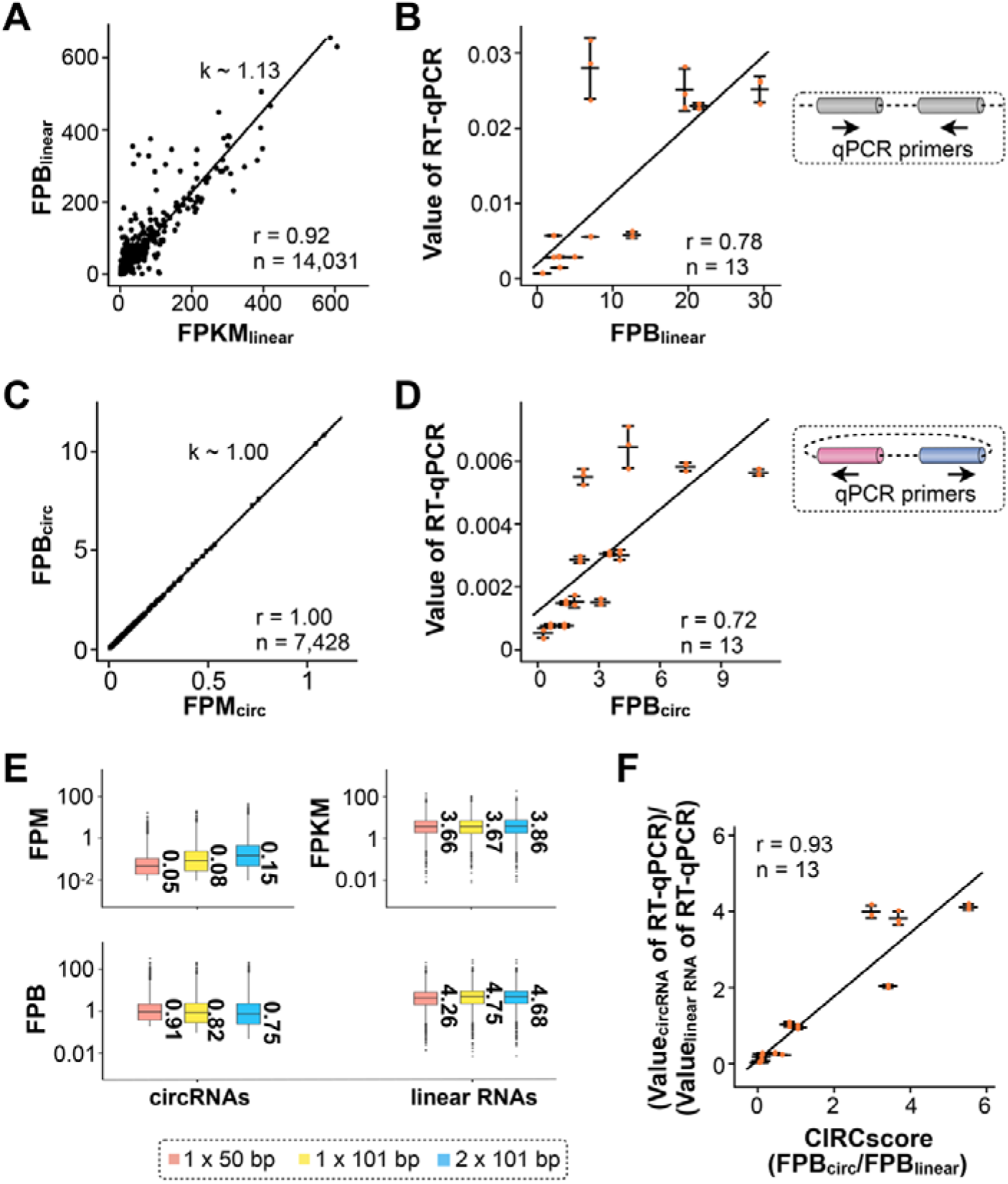
Comparison of FPB with other quantification statistics. (**A**) Comparison of FPB_linear_ and FPKM_linear_. FPB_linear_ is highly correlated with FPKM_linear_ (Pearson correlation coefficient (PCC) = 0.92) in PA1 cells, under the condition of both FPB_linear_ and FPKM_linear_ > 0. The slope (k) was calculated from linear regression by the lm function in R. (**B**) FPB_linear_ is highly correlated with linear RNA expression measured by RT-qPCR. Expression of thirteen linear RNAs (Supplementary Table S3) was measured by RT-qPCR, and highly correlated with FPB_linear_ values obtained from PA1 ribo– RNA-seq (PCC = 0.78). Right, qPCR primers are convergently spanned linear RNA exons. Means ± s.d. were from three independent experiments. (**C**) Comparison of FPB_circ_ and FPM_circ_. FPB_circ_ is highly correlated with FPM_circ_ (PCC = 1.00) in PA1 cells. The slope (k) was calculated from linear regression by the lm function in R. (**D**) FPB_circ_ is highly correlated with circRNA expression measured by RT-qPCR. Expression of thirteen circRNAs (Supplementary Table S3) was measured by RT-qPCR, and highly correlated with FPB_circ_ values obtained from PA1 ribo– RNA-seq data (PCC = 0.72). Right, qPCR primers are divergently spanned circRNA exons. Means ± s.d. were from three independent experiments. (**E**) FPB is resistant to the changes of sequencing lengths and strategies. Two virtual RNA-seq datasets were constructed from original 2 × 101 bp cortex ribo– RNA-seq dataset to mimic different sequencing lengths and strategies, including 1 × 101 bp (extract the left part of reads from the paired-end dataset) and 1 × 50 bp (extract first 50 bp read sequence from left part of reads). All the three RNA-seq datasets were used for circular and linear RNA quantification to obtain related FPM, FPB and/or FPKM values. Unlike FPM, FPB largely remained unchanged with different sequencing lengths and strategies. Note, FPKM was set as a control that was also not largely altered by the changes of sequencing lengths and strategies. (**F**) CIRCscore is highly correlated with the value of circular *vs* linear RNA expression measured by RT-qPCR (PCC = 0.93). Means ± s.d. were from three independent experiments.

### Comparison of FPB with FPM for circRNA quantification

As expected, circRNA expression, when quantitated by FPB_circ_, is highly correlated with that by FPM_circ_ (Figure 2C). Experimentally, FPB_circ_ is also highly correlated with the expression of thirteen examined circRNAs measured by RT-qPCR in PA1 cells (Figure 2D, Supplementary Table S3). The expression of these thirteen circRNAs ranges from ∼ 1 to 10 FPB (Figure 2D), and their cognate linear RNAs are evaluated above (Figure 2B).

Importantly, compared to commonly-used FPM, FPB is resistant to different sequencing lengths and strategies, such as 1×50 *vs* 1×100 or single- *vs* paired-end RNA-seq datasets (Figure. 2E). These results are in reasonable agreement with the definitions of FPB and FPM. For example, 1 FPB is equivalent to 0.1 FPM for 1 × 100 bp single-end RNA-seq datasets (Supplementary Figure S5A) and to 0.2 FPM for 2 × 100 bp paired-end RNA-seq datasets (Supplementary Figure S5B). In this scenario, FPB can be used directly for cross-sample comparison regardless of different sequencing lengths and strategies.

### Evaluation of relative circRNA expression by CIRCscore

Different from unscaled and non-comparable values of FPKM_linear_ (for linear RNA expression) and FPM_circ_ (for circRNA expression), FPB_linear_ for linear RNA measurement is comparable to FPB_circ_ for circRNA measurement. We divide FPB_circ_ by FPB_linear_ to obtain CIRCscore values, by which expression levels of circular and linear RNAs are directly compared in a genome-wide manner. Importantly, the CIRCscore was highly correlated with the experimental comparison of circular *vs* linear RNA expression measured by RT-qPCR in thirteen examined gene loci from PA1 cells (Figure 2F and Supplementary Table S3), confirming the notion that CIRCscore provides an additional parameter to evaluate circRNA expression normalized by their cognate linear RNA expression background.

Since circRNAs are generally co-expressed with their cognate linear RNAs and that sequences of circRNAs largely overlap with those of linear RNAs, the advantage of using CIRCscore to quantitate circRNA expression is that it normalizes their expression to the linear RNA expression background. As shown for in the PA1 cell line, among those with FPB_circ_ ≥ 1, some circRNAs with high FPB values exhibit low CIRCscore values (Figure 3A, blue), which is likely due to the high expression of their cognate linear RNAs (Figure 3B). However, other circRNAs with comparable FPB values have relatively high CIRCscores (Figure 3A, red), as their cognate linear RNAs are expressed at low levels (Figure 3C). This observation suggests variable expression patterns of circular and their cognate linear RNAs in different genomic loci.

**Figure 3.**
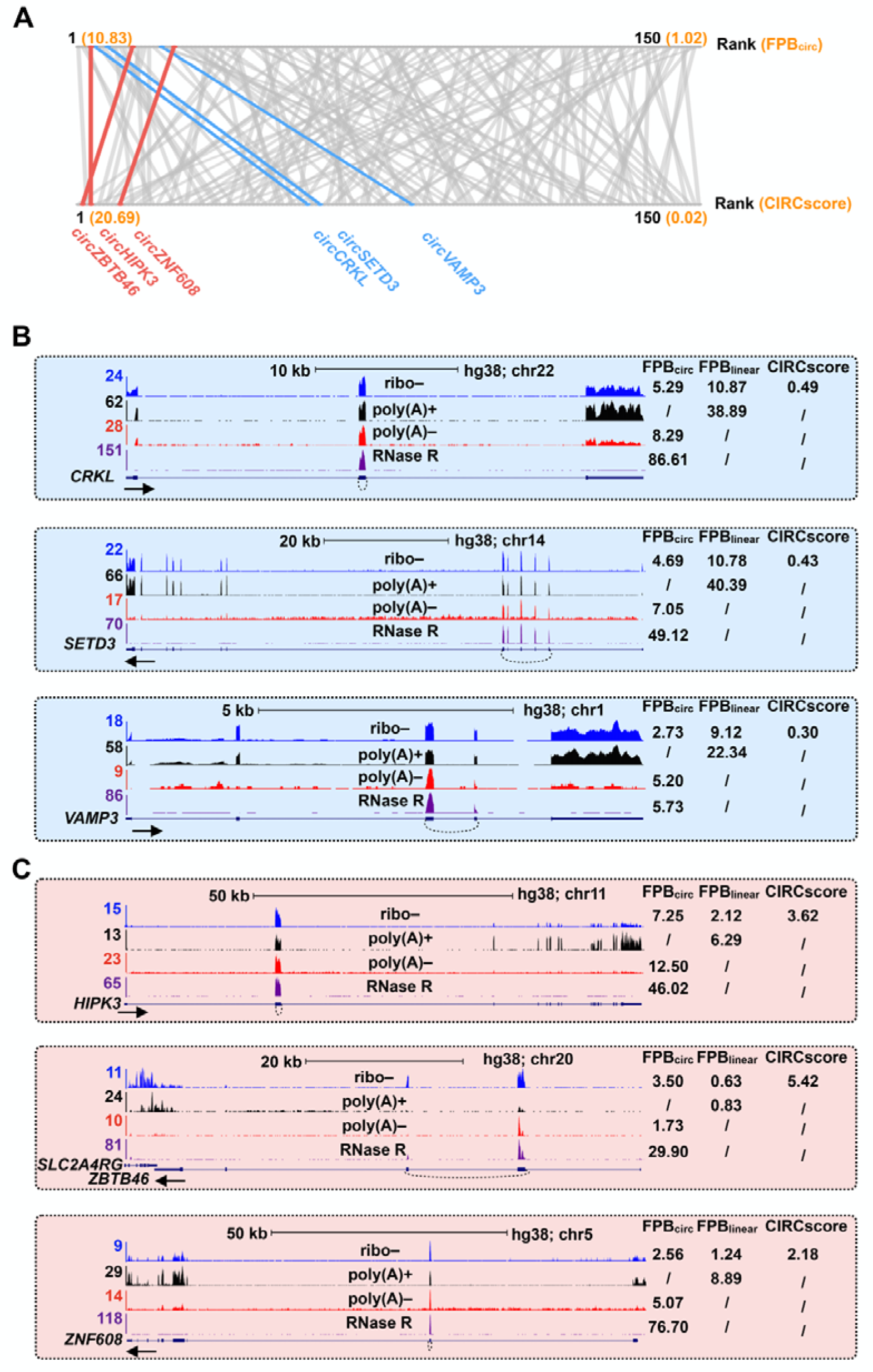
Comparison of circRNA quantification by FPB and CIRCscore. (**A**) Difference of circRNAs quantitated by FPB_circ_ or CIRCscore in PA1 cells. About 150 circRNAs with FPB_circ_ ≥ 1 are identified in PA1 cells from published ribo– RNA-seq (17,18). Some circRNAs with high FPB_circ_ values exhibit high CIRCscore values due to the low background of cognate linear RNA expression (Red), while some others show low CIRCscore values due to the high background of cognate linear RNA expression (Blue). (**B**) Three highly-expressed circRNAs, *circCRKL, circSETD3* and *circVAMP3*, are co-expressed with their cognate linear RNAs at high levels, indicated by relatively low CIRCscores. (**C**) Three highly-expressed circRNAs, *circHIPK3, circZBTB46* and *circZNF608* are co-expressed with their cognate linear RNAs at low levels, indicated by relatively high CIRCscores.

We further applied CLEAR to evaluate circRNAs in twelve additional human tissues with both FPB and CIRCscore values (Figure 4A and Supplementary Table S4). Consistent with previous findings (26), circRNAs are more abundant in brain samples than in non-brain tissues. Among all six examined brain samples, circRNAs are more enriched in the cortex, occipital and diencephalon, but less in the cerebellum, when evaluated by both FPB (Figure 4A, left) and CIRCscore (Figure 4B, right) values. In six non-brain tissues, circRNAs are enriched in the heart and thyroid at a comparable level as in the cerebellum. About 10%-20% of circRNAs with FPB_circ_ ≥1 are expressed at a comparable or even higher level than their cognate linear RNAs, indicated by CIRCscore ≥1 (Figure 4A, right), such as in gene loci for *circTPTE2P5* and *circPHF7* (Figure 4B). Taken together, the identification of highly-expressed circRNAs with high FPB_circ_ and CIRCscore values reveals that some gene loci are particularly favorable for circRNA production (Supplementary Table S4), and such circRNAs are candidates for subsequent functional studies.

**Figure 4.**
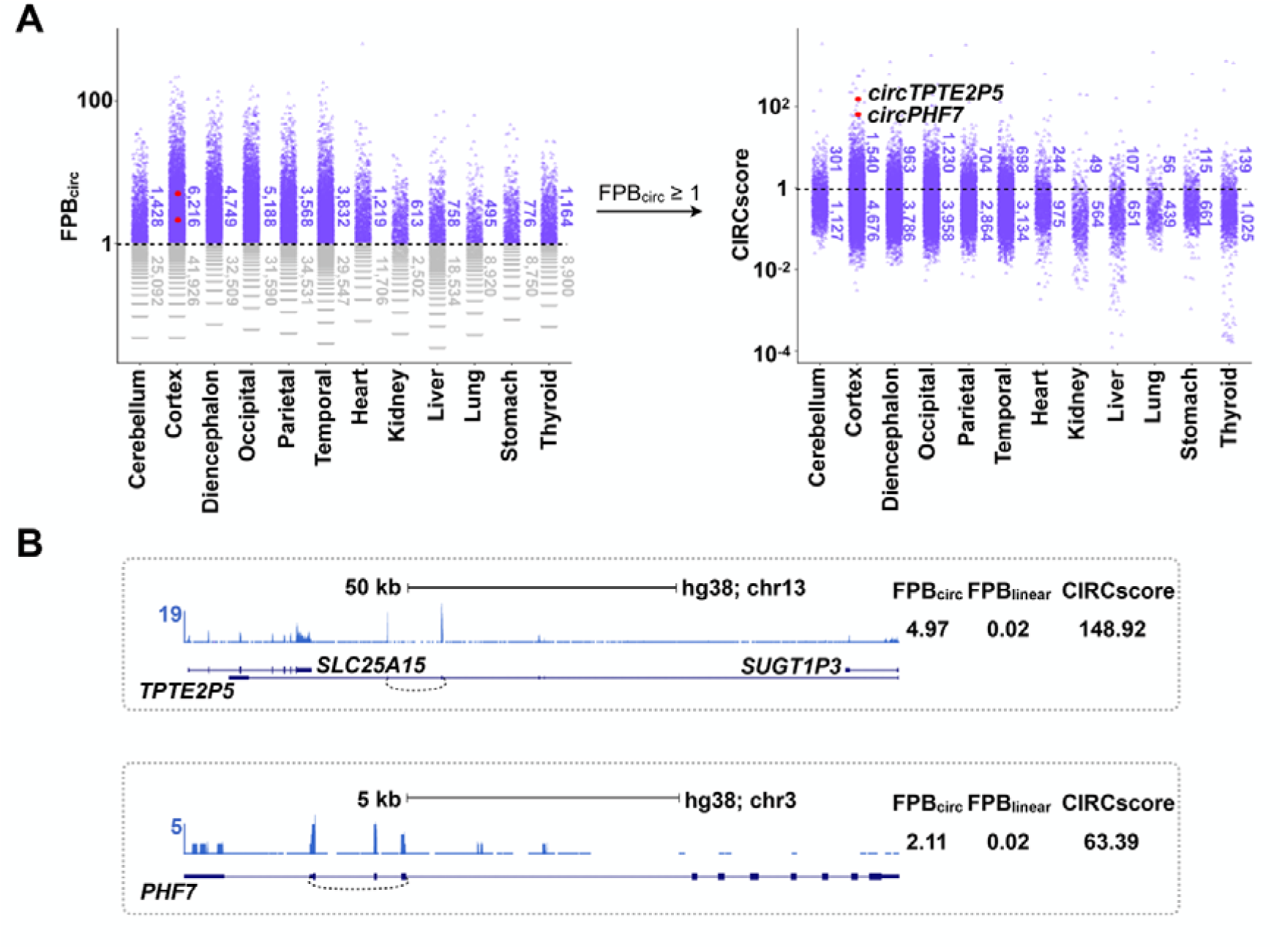
Quantification of circRNAs among twelve human tissue samples. (**A**) Quantification of circRNAs by the CLEAR pipeline. CircRNAs in twelve ENCODE human tissues are quantitated by FPB_circ_ (left), and those with FPB_circ_ ≥ 1 (purple) are further evaluated by CIRCscore (Right). Two represented circRNAs in cortex tissue with fair FPB but high CIRCscore values are highlighted in red. (**B**) Visualization of *circTPTE2P5* and *circPHF7* in ENCODE human cortex sample. Of Note, *circTPTE2P5* and *circPHF7* are co-expressed with their cognate linear RNAs at low levels, indicated by high CIRCscores.

### CIRCscore reduces individual differences

Different to FPB, using CIRCscore to evaluate circRNA expression can reduce individual differences that are caused by RNA-seq samples themselves. For example, compared to paired control samples, circRNA expression evaluated by the FPB_circ_ value is inconsistent in a batch of 20 human hepatocellular carcinoma (HCC) samples (GSE77509) (20). Some HCC samples appear to have generally low circRNA expression; while others, such as samples #11 and #16, appear to have significantly high circRNA expression (Supplementary Figure S6A). Consequently, it is hard to distinguish circRNA expression differences between HCC and their paired control samples using FPB_circ_ in these samples (Figure 5A, *P* = 0.99). Strikingly, it is clearly shown that circRNAs are generally lowly expressed in almost all HCC samples when CIRCscore is used to normalize circRNA expression with cognate linear RNA background (Figure 5B, *P* = 3.59 × 10^−5^ and Supplementary Figure S6B). These results suggest that it is important to take cognate linear RNA expression into consideration for circRNA quantification, which can be achieved by the CLEAR pipeline in a genome-wide manner. Taken together, quantification of circRNA expression by CIRCscore helps to eliminate individual differences among paired comparisons, and it can therefore be used to decipher the trend of circRNA expression changes during different conditions and diseases across RNA-seq datasets.

**Figure 5.**
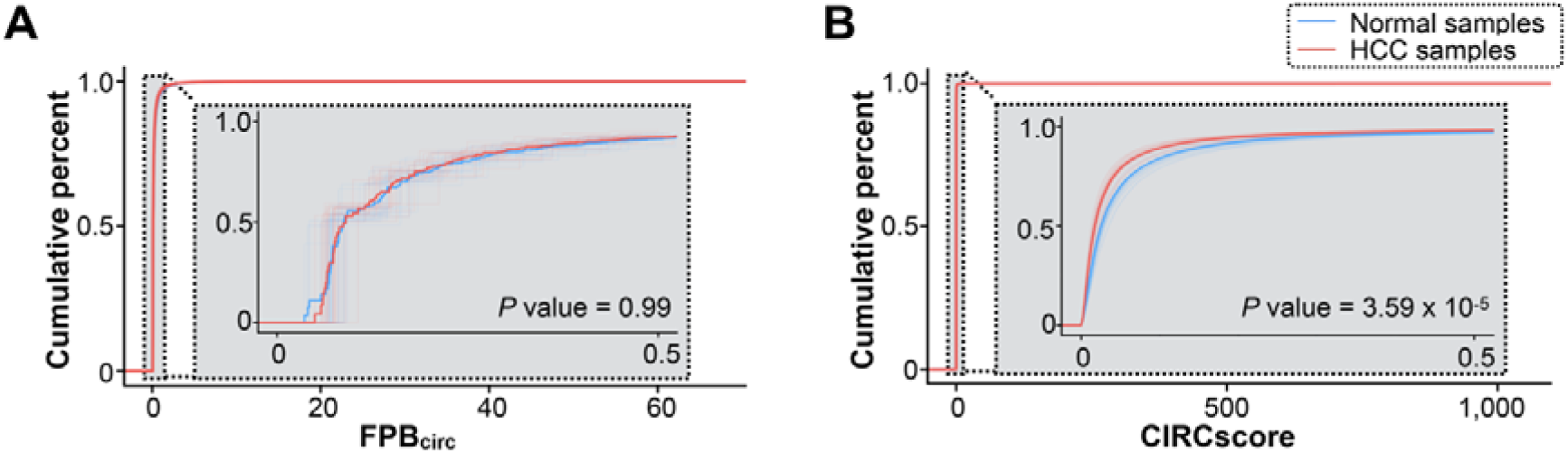
Removal of sample/batch effects with CIRCscore quantification. (**A**) Cumulative distribution and comparison of circRNAs in 20 paired human hepatocellular carcinoma (HCC) and normal control samples by FPB_circ_. Thick blue and red line represent the mixture distribution of FPB_circ_ from 20 normal or HCC samples, respectively. *P* value for statistical significance of difference between two distributions (normal *vs* HCC) was calculated using two-tailed unpaired Student’s *t*-test. (**B**) Cumulative distribution and comparison of circRNAs in 20 paired HCC and normal control samples by CIRCscore. Thick blue and red line represent the mixture distribution of CIRCscores from 20 normal or HCC sample, respectively. *P* value for statistical significance of difference between two distributions (normal *vs* HCC) was calculated using two-tailed unpaired Student’s t-test.

## DISCUSSION

Recently, circRNAs have been widely detected in examined cell lines and tissues by deep sequencing of non-polyadenylated RNAs together with specific computational pipelines for detecting RNA-seq reads/fragments mapped to BSJ sites (17,26,27). Due to distinct strategies for circular or linear RNA quantification (Figure 1A), computational pipelines for direct circular and linear RNA expression comparison from RNA-seq datasets have until now been unavailable. In this study, we have developed CLEAR by applying normalized RNA-seq fragments solely mapped to BSJ or canonical SJ sites individually for circular (FPB_circ_) or cognate linear (FPB_linear_) RNA quantification (Figure 1B).

The CLEAR pipeline has at least two advantages in circRNA studies. First, the FPB values are highly correlated with canonical FPKMs for linear RNAs and FPMs for circRNAs (Figure 2), and unlikely affected by RNA-seq strategies, which makes cross-sample comparisons feasible. Second, direct comparison of circular and cognate linear RNAs with the CIRCscore not only precisely quantitates relative circRNA expression normalized by linear RNA expression background (Figure 3 and 4), but also eliminates possible errors/fluctuations caused by sample preparation/sequencing differences (Figure 5), which reduces inaccuracies for circRNA quantification and subsequent cross-sample comparison. Thus, CLEAR has the potential to allow one to identify highly expressed circRNAs in different biological settings for subsequent functional studies. This is important, because so far individual circRNA functions remain to be explored as it is hard even to identify those with high expression levels in the context of interest.

It is worthwhile noting that different RNA sequencing strategies have been applied to profile circRNAs, including ribo-, poly(A)–/ribo– and RNase R-treated RNA-seq datasets (Figure 3). Different from poly(A)+ RNA-seq datasets that are used to detect polyadenylated cognate linear RNAs, all three types of non-polyadenylated RNA-seq can be used to determine circRNA expression by FPB. However, only ribo-RNA-seq datasets that profile both polyadenylated linear and non-polyadenylated circular RNAs in parallel are suitable for direct circular and linear RNA expression comparison by CIRCscore (Figure 3). In contrast, in poly(A)–/ribo– and RNase R-treated RNA-seq datasets, polyadenylated linear RNAs are largely depleted, which is unsuitable for accurate linear RNA quantification and subsequent CIRCscore evaluation.

Taken together, the CLEAR pipeline provides a comprehensive way to quantitatively evaluate circRNA expression across samples and to identify highly expressed circRNAs with low linear RNA expression background.

## AVAILABILITY

The CLEAR pipeline and its application can be downloaded from https://github.com/YangLab/CLEAR.

## SUPPLEMENTARY DATA

Supplementary Data are available online.

## ACKNOWLEDGEMENT

We like to acknowledge lab members for testing the CLEAR toolkit.

