## Supplementary data for "A CLEAR pipeline for direct comparison of circular and linear RNA expression"

**Supplementary figures and figure legends**

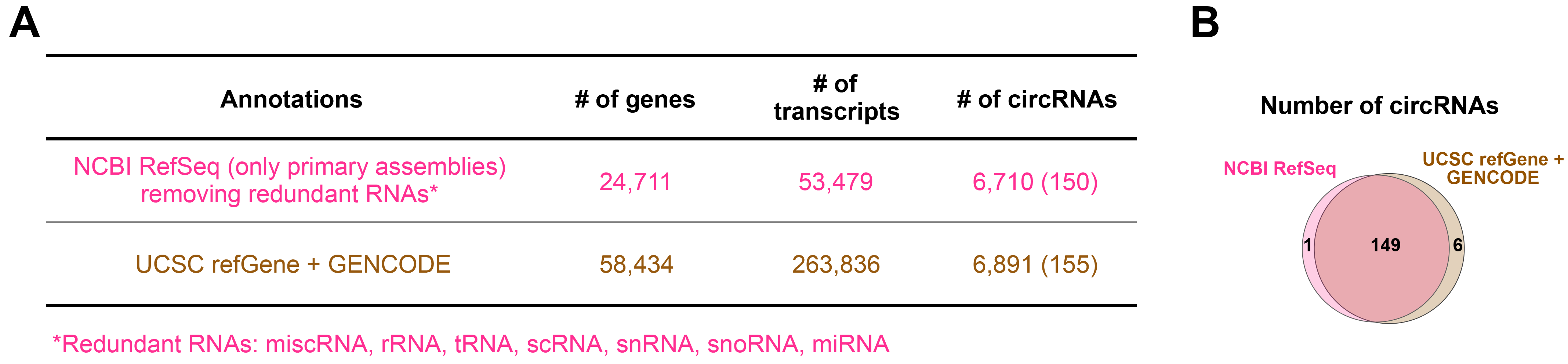

**Supplementary Figure S1. Gene annotation used for circular and linear RNA expression**

**analysis.**

(**A**) Two commonly used gene annotations, NCBI RefSeq and the combination of UCSC refGene together with GENCODE annotations, were compared in this study. Although different genes and transcripts were obtained from two gene annotations, similar circRNAs were predicted with the CLEAR pipeline. Thus, the NCBI RefSeq annotation was used in this study for circular and linear RNA expression analysis after removing redundant RNAs.

(**B**) Almost completely overlap of circRNAs predicted by the CLEAR pipeline with two different gene annotations.

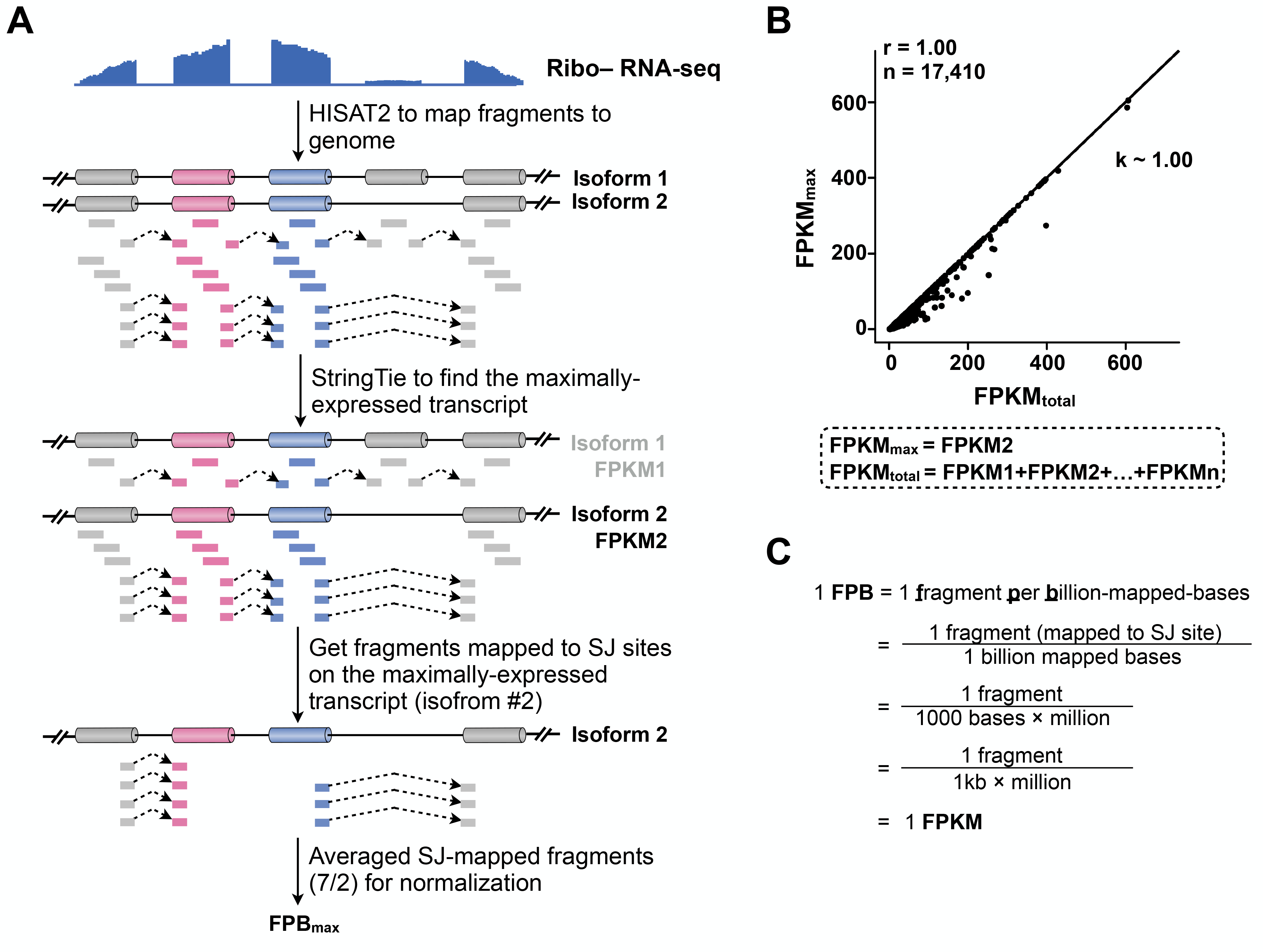

**Supplementary Figure S2. Selection of maximally-expressed linear transcripts for analyses.**

(**A**) Schematic drawing to show the selection of maximally-expressed linear transcripts for analyses. StringTie was used to find and calculate the expression of different transcripts from given gene loci in CLEAR pipeline. See Methods for details.

(**B**) Expression level of maximally-expressed linear transcripts can be used to represent gene expression of given loci. FPKM_max_ of maximally-expressed linear transcripts of given loci is highly correlated with FPKM_total_ of all transcripts of given loci (Pearson correlation coefficient (PCC) = 1.00). PA1 ribo– RNA-seq dataset was used for the comparison. The slope (k) was calculated from linear regression by lm function in R.

(**C**) A formula to show that the value of FPB is theoretically equivalent to the value of FPKM.

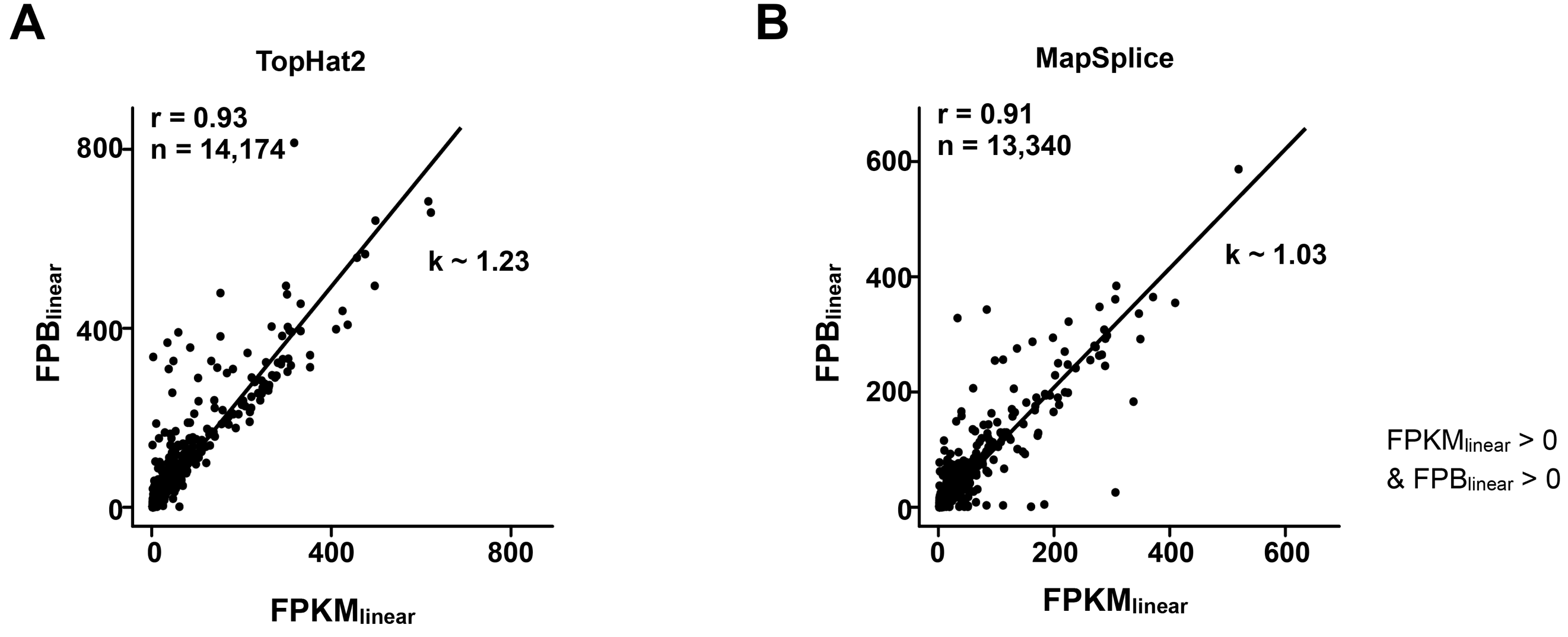

**Supplementary Figure S3. High correlation between FPB and FPKM for linear RNA quantification with different aligners.**

(**A**) FPB_linear_ is highly correlated with FPKM_linear_ (PCC = 0.93) in PA1 ribo– RNA-seq data by using TopHat2 for mapping, under the condition of both FPB_linear_ > 0 and FPKM_linear_ > 0. The slope (k) was calculated from linear regression by lm function in R.

(**B**) FPB_linear_ is highly correlated with FPKM_linear_ (PCC = 0.91) in PA1 ribo– RNA-seq data by using MapSplice for mapping, under the condition of both FPB_linear_ > 0 and FPKM_linear_ > 0. The slope (k) was calculated from linear regression by lm function in R.

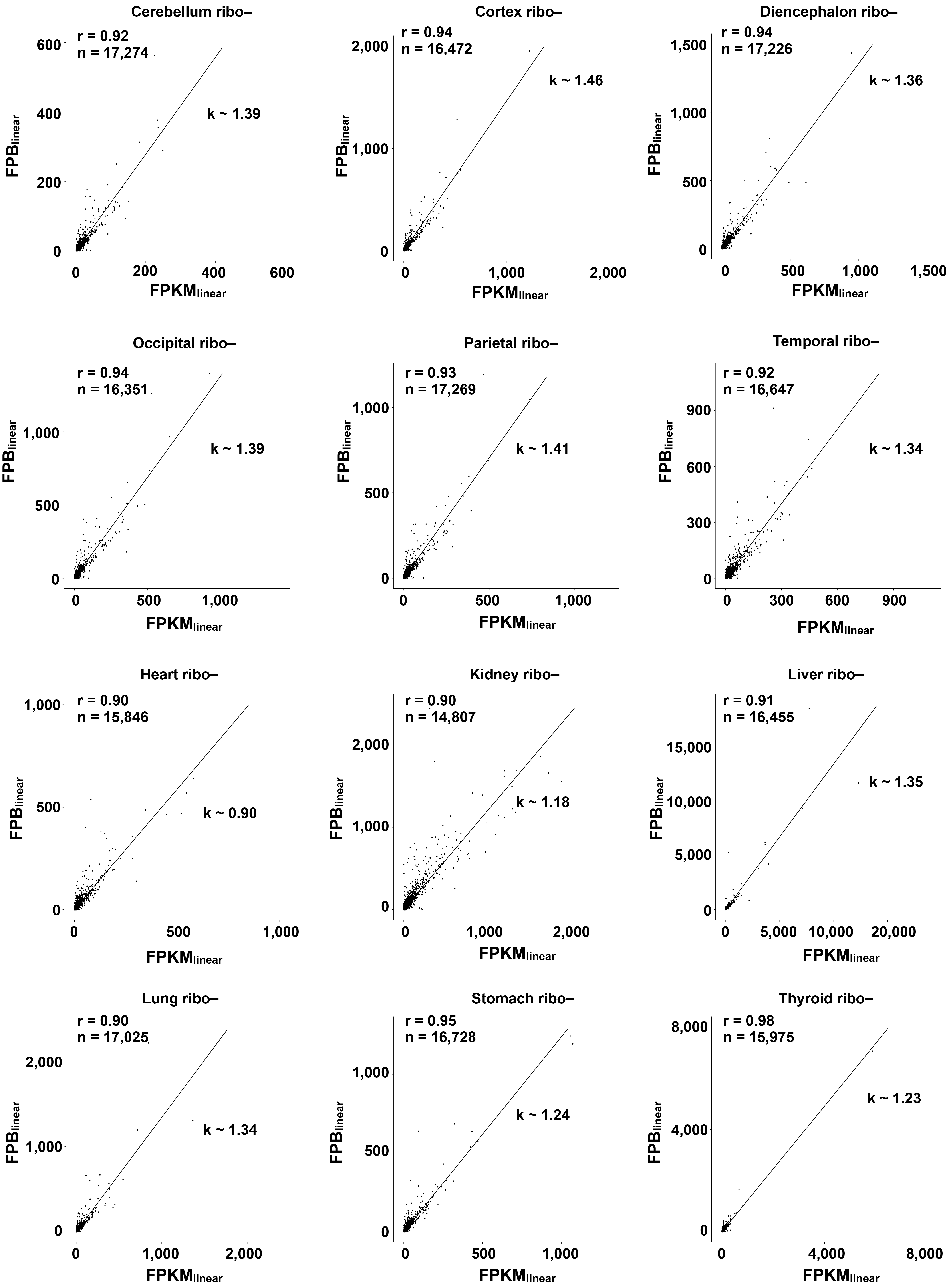

**Supplementary Figure S4. High correlation between FPB and FPKM for linear RNA quantification.**

FPB_linear_ is highly correlated with FPKM_linear_ in all examined 12 ribo– samples from different human tissues (PCC were between 0.90 and 0.98), under the condition of both FPB_linear_ > 0 and FPKM_linear_ > 0. The slope (k) was calculated from linear regression by the lm function in R.

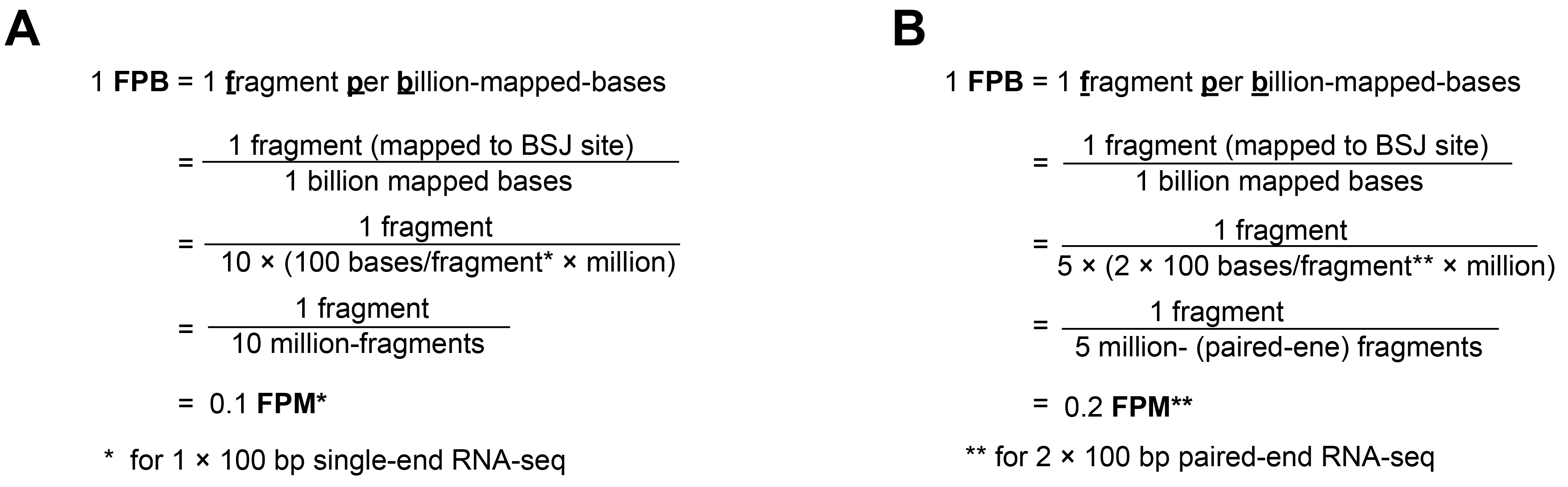

**Supplementary Figure S5. Comparison of FPB and FPM.**

(**A**) A formula to show that the value of 1 FPB is theoretically equivalent to the value of 0.1 FPM by 1 × 100 bp single-end RNA-seq.

(**B**) A formula to show that the value of 1 FPB is theoretically equivalent to the value of 0.2 FPM by 2 × 100 bp paired-end RNA-seq.

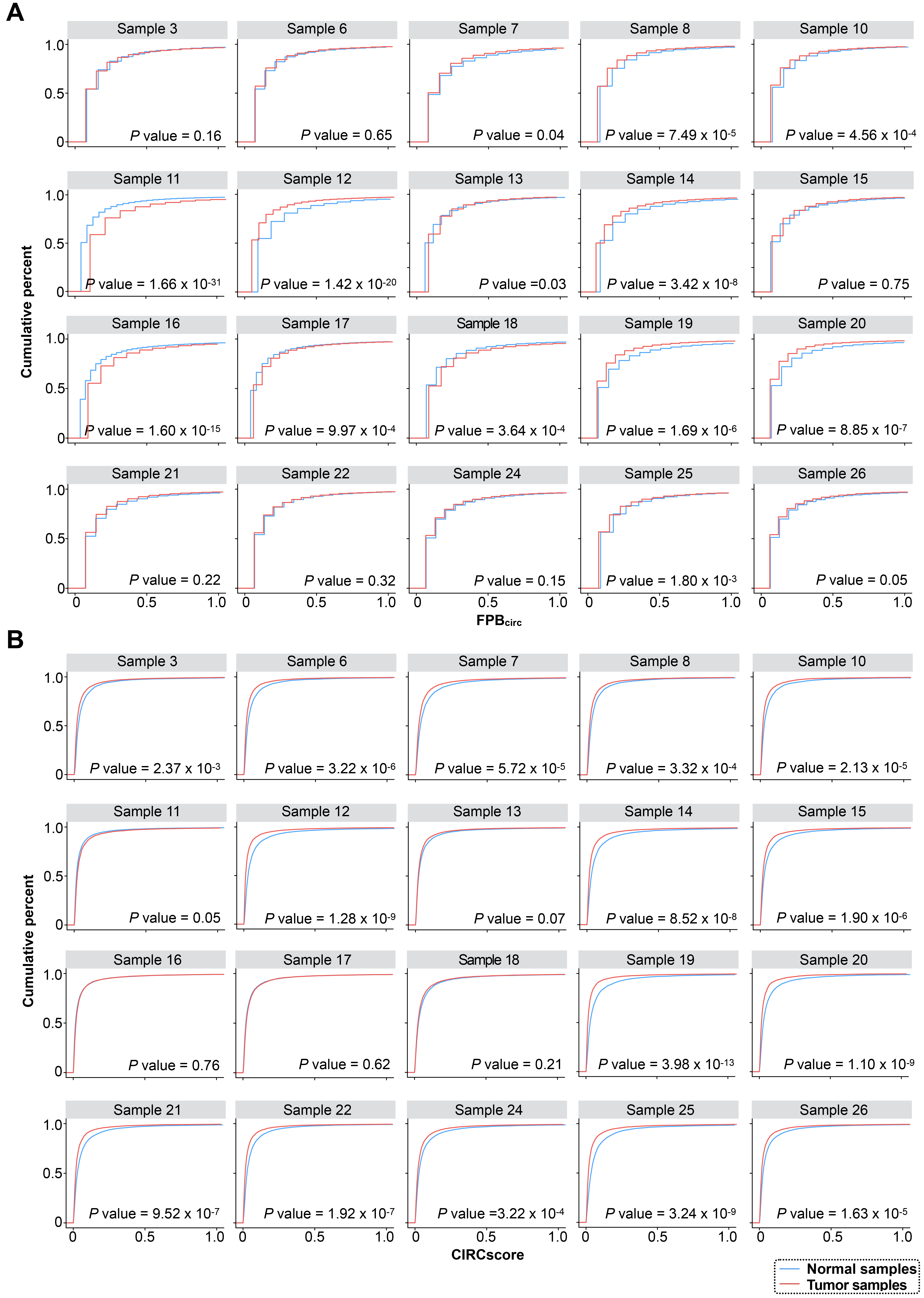

**Supplementary Figure S6. Comparison of using FPB_circ_ or CIRCscore to evaluate circRNA expression in paired normal and tumor samples.**

(**A**) Cumulative distribution of circRNA expression by FPB_circ_. Blue or red line represents FPB_circ_ distribution of each paired normal and HCC sample, respectively.

(**B**) Cumulative distribution of relative circRNA expression by CIRCscore. Blue or red line represents CIRCscore distribution of each paired normal and HCC samples, respectively.

**Supplementary tables and table legends**

| **Primer sequences used in RT-qPCR for comparing with FPB and CIRCscore** | |
| --- | --- |
| **CORO1C-C-F1** | CTCAGAGAGGGATGGGTTACA |
| **CORO1C-C-R1** | ACCCCATTGCTAGTGTCCAT |
| **CORO1C-L-F1** | CTGCACAGCTTCCAAAGACA |
| **CORO1C-L-R1** | TGGGTCTTGCTCCTTCATGT |
| **FKBP8-C-F1** | CGGCCATGGTCACTGCTG |
| **FKBP8-C-R1** | ACCAGCGTCTTCTTCCTCAA |
| **FKBP8-L-F1** | CGCCAACTCCTACGACCTC |
| **FKBP8-L-R1** | GACACTTCACCTTCAACTGCA |
| **KIAA0368-C-F1** | AGAAATTCTTCAAGATTTGG |
| **KIAA0368-C-R1** | TTGAAATACCACTGTCTCTCC |
| **KIAA0368-L-F1** | GCCTGGCAAATGACTGAAGA |
| **KIAA0368-L-R1** | GTGGGTTGGGGCTGATGATA |
| **SMO-C-F1** | TCATCGTGGGAGGCTACTTC |
| **SMO-C-R1** | GTTGTCTGTCCGAACCAAGG |
| **SMO-L-F1** | CTGCCAGCAAGATCAACGAG |
| **SMO-L-R1** | GGCAGCTGAAGGTAATGAGC |
| **ARHGAP-C-F1** | CCGCAGGAGAAGGCTCTGAA |
| **ARHGAP-C-R1** | GGAGATCTACACCAGTATCA |
| **ARHGAP-L-F1** | GTTGGACTCGGGATGCAAG |
| **ARHGAP-L-R1** | ACTGACTATCTGACTGGCTGT |
| **HIPK3-C-F1** | GTCGGCCAGTCATGTATCAA |
| **HIPK3-C-R1** | ACCAAGACTTGTGAGGCCAT |
| **HIPK3-L-F1** | TGGGATGTGTGATTGCAGAA |
| **HIPK3-L-R1** | AACTGTTCTCCTGGCAAACC |
| **CAMSAP1-C-F1** | GAGGATGCCATGGTGTTCTG |
| **CAMSAP1-C-R1** | TAACAGGCGGCTTAATGTGC |
| **CAMSAP1-L-F1** | GAAAGTCCAGCTCATCAAAAG |
| **CAMSAP1-L-R1** | TCCAACAACGGGAAGTAGGG |
| **ZBTB46-C-F1** | CTCCCTGCTGTTCGAGTACC |
| **ZBTB46-C-R1** | CATTTGGAAGCCCTGGTGTC |
| **ZBTB46-L-F1** | CCTACCCCTGCGAGATCTG |
| **zBTB46-L-R1** | ACTTCTTGTCCTTGCTGTGG |
| **CAPRIN1-C-F1** | CATGCCTCCATTCACCTGTG |
| **CAPRIN1-C-R1** | AAGCCTTTCCCCTTTGTTCA |
| **CAPRIN1-L-F1** | ATGGGCTGTGTGAGGAAGAA |
| **CAPRIN1-L-R1** | CAGTGTACTCTTCTGCTGGC |
| **CDK8-C-F1** | CCTCCTACCACTACCTCAGG |
| **CDK8-C-R1** | GGTCCTGCATAGCCTGTTCT |
| **CDK8-L-F1** | CCAACTGCAGCCTTATCAAGT |
| **CDK8-L-R1** | GGTCCATGGTAAGCAGCTTC |
| **MGA-C-F1** | GTGAAGTTCCTCAATTGAAGCA |
| **MGA-C-R1** | GCTGTTTCTCCTCCATGATTTCA |
| **MGA-L-F1** | TGGAAGATGTGGATGGTGTTC |
| **MGA-L-R1** | TTGTGGTTGATGCCTGGAGA |
| **FAM13B-C-F1** | TCAAAACCTGTGGCTAGCAC |
| **FAM13B-C-R1** | GAGGCTGGTAGGATGCTGAT |
| **FAM13B-L-F1** | TGGCTGGACTTCTGGAAAAC |
| **FAM13B-L-R1** | TGTTCCAGCTTTTCATCTTCCTC |
| **PLEKHM3-C-F1** | GGAACGAGGACTCACTGCT |
| **PLEKHM3-C-R1** | CAGCTTTGCCAGTAACTGTCA |
| **PLEKHM3-L-F1** | CTTTCTCATTCCAGCACGCA |
| **PlEKHM3-L-R1** | CTCCAGAAACTCCTTGGCCT |

**Supplementary Table S1. List of primers used in this study.**

Primer sequences used in RT-qPCR for comparing with FPB and CIRCscore are listed.

| **Species** | **Cell line/tissues** | **Type** | **Layout** | **Read length (bp)** | **SRA ID** | **# of pairs** |
| --- | --- | --- | --- | --- | --- | --- |
| Human | PA1 | ribo– | Single-end | 100 | SRR2483517,  SRR2483518 | 125,735,340 |
|  |  | poly(A)+ |  |  | SRR2976714 | 52,014,532 |
|  |  | poly(A)– |  |  | SRR2976715 | 87,675,979 |
|  |  | RNase R |  |  | SRR2976716 | 69,931,871 |
|  | Cerebellum | ribo– | Paired-end | 101 | SRR319242 | 121,997,943 |
|  | Cortex |  |  |  | SRR3192424 | 114,797,948 |
|  | Diencephalon |  |  |  | SRR3192431 | 78,265,226 |
|  | Occipital |  |  |  | SRR3192443 | 90,751,416 |
|  | Parietal |  |  |  | SRR3192445 | 102,321,119 |
|  | Temporal |  |  |  | SRR3192463 | 142,047,587 |
|  | Heart |  |  |  | SRR3192433 | 77,191,913 |
|  | Kidney |  |  |  | SRR3192604 | 106,249,338 |
|  | Liver |  |  |  | SRR3192418 | 166,369,413 |
|  | Lung |  |  |  | SRR3192441 | 115,978,515 |
|  | Stomach |  |  |  | SRR3192449 | 63,403,860 |
|  | Thyroid |  |  |  | SRR4422152,  SRR4422153 | 81,957,307 |

**Supplementary Table S2. Summary of ribo− RNA-seq datasets from various human cell line/tissues used in this study.**

Information including species, cell line/tissues, RNA-seq strategies, SRA IDs and total sequencing read counts are listed in the table.

| **Gene locus** | **Location of circRNA** | **FPB_circ_** | **Relative Ct value of circRNA (to actin)** | **FPB_linear_** | **Relative Ct value of linear RNA (to actin)** | **CIRCscore** | **Ratio of circRNA/linear RNA by RT-qPCR** |
| --- | --- | --- | --- | --- | --- | --- | --- |
| CORO1C | chr12:108652271-108654410 | 1.28 | 10.32 | 29.44 | 5.26 | 0.04 | 0.03 |
|  |  |  | 10.46 |  | 5.43 |  | 0.03 |
|  |  |  | 10.27 |  | 5.25 |  | 0.03 |
| FKBP8 | chr19:18539370-18539720 | 1.79 | 9.47 | 21.35 | 5.43 | 0.08 | 0.06 |
|  |  |  | 9.45 |  | 5.43 |  | 0.06 |
|  |  |  | 9.17 |  | 5.47 |  | 0.08 |
| KIAA0368 | chr9:111386376-111391824 | 1.36 | 9.43 | 12.52 | 7.31 | 0.11 | 0.23 |
|  |  |  | 9.33 |  | 7.49 |  | 0.28 |
|  |  |  | 9.41 |  | 7.47 |  | 0.26 |
| SMO | chr7:129205202-129206587 | 3.07 | 9.41 | 7.03 | 7.48 | 0.44 | 0.26 |
|  |  |  | 9.27 |  | 7.50 |  | 0.29 |
|  |  |  | 9.44 |  | 7.47 |  | 0.25 |
| ARHGAP12 | chr10:31908171-31910563 | 4.01 | 8.35 | 4.90 | 8.42 | 0.82 | 1.05 |
|  |  |  | 8.46 |  | 8.42 |  | 0.97 |
|  |  |  | 8.31 |  | 8.44 |  | 1.09 |
| HIPK3 | chr11:33286412-33287511 | 7.25 | 7.41 | 2.12 | 8.45 | 3.41 | 2.05 |
|  |  |  | 7.46 |  | 8.45 |  | 1.99 |
|  |  |  | 7.40 |  | 8.45 |  | 2.07 |
| CAMSAP1 | chr9:135881632-135883078 | 10.83 | 7.44 | 2.93 | 9.44 | 3.68 | 4.02 |
|  |  |  | 7.49 |  | 9.37 |  | 3.68 |
|  |  |  | 7.48 |  | 9.39 |  | 3.76 |
| ZBTB46 | chr20:63775677-63790790 | 3.5 | 8.33 | 0.63 | 10.35 | 5.52 | 4.05 |
|  |  |  | 8.38 |  | 10.45 |  | 4.20 |
|  |  |  | 8.36 |  | 10.39 |  | 4.09 |
| CAPRIN1 | chr11:34071725-34076642 | 0.6 | 10.47 | 19.49 | 5.46 | 0.03 | 0.03 |
|  |  |  | 10.33 |  | 5.35 |  | 0.03 |
|  |  |  | 10.26 |  | 5.15 |  | 0.03 |
| CDK8 | chr13:26400452-26401624 | 0.26 | 10.66 | 2.84 | 8.41 | 0.09 | 0.21 |
|  |  |  | 11.46 |  | 8.45 |  | 0.12 |
|  |  |  | 10.64 |  | 8.38 |  | 0.21 |
| MGA | chr15:41668827-41669958 | 4.43 | 7.14 | 7.00 | 4.98 | 0.63 | 0.22 |
|  |  |  | 7.26 |  | 5.13 |  | 0.23 |
|  |  |  | 7.44 |  | 5.40 |  | 0.24 |
| FAM13B | chr5:137985256-137988315 | 2.22 | 7.45 | 2.13 | 7.47 | 1.04 | 1.01 |
|  |  |  | 7.58 |  | 7.43 |  | 0.90 |
|  |  |  | 7.49 |  | 7.43 |  | 0.96 |
| PLEKHM3 | chr2:207976650-207977586 | 2.05 | 8.39 | 0.68 | 10.46 | 2.97 | 4.18 |
|  |  |  | 8.46 |  | 10.42 |  | 3.89 |
|  |  |  | 8.49 |  | 10.45 |  | 3.90 |

**Supplementary Table S3. Summary of validated circRNAs and their linear cognate RNAs.**

Information including gene locus, genomic location, FPB and relative Ct value (from three independent experiments) of thirteen circRNAs and their linear cognate RNAs validated in this study are listed. CIRCscore and ratios of circRNA/linear RNA performed by RT-qPCR (from three independent experiments) are listed.

**Supplementary Table S4. Quantification of circRNAs by both FPB and CIRCscore from ribo− RNA-seq datasets in twelve human tissues.**

Information including genomic location, strand, gene symbol, transcript id, included exons, flanking introns, FPB_circ_, FPB_linear_ and CIRCscore from twelve human tissues including cerebellum, cortex, diencephalon, occipital, parietal, temporal, heart, kidney, liver, lung, stomach and thyroid are listed respectively.
